# Conditional repeatability and the variance explained by reaction norm variation in random slope models

**DOI:** 10.1101/2020.03.11.987073

**Authors:** Holger Schielzeth, Shinichi Nakagawa

## Abstract

Individuals differ in average phenotypes and in sensitivity to environmental variation. Such context-sensitivity can be modelled as random-slope variation. Random-slope variation implies that the proportion of between-individual variation varies across the range of a covariate (environment/context/time/age) and has thus been called ‘conditional’ repeatability. We propose to put conditional repeatabilities in perspective of the total phenotypic variance and suggest a way of standardization using the random-slope coefficient of determination 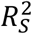. Furthermore, we illustrate that the marginalized repeatability *R_mar_* averaged across an environmental gradient offers a biologically relevant description of between-individual variation. We provide simple equations for calculating key descriptors of conditional repeatabilities, clarify the difference between random-intercept variation and average between-individual variation and make recommendations for comprehensive reporting. While we introduce the concept with individual variation in mind, the framework is equally applicable to other type of between-group/cluster variation that varies across some (environmental) gradient.

## Introduction

Repeatabilities *R* and coefficients of variation *R*^2^ allow a decomposition of the sources of biological variance in some response, feature or trait of interest. Repeatabilities (also known as intra-class correlations, ICC) are concerned with a decomposition of random-effect variances (Nakagawa and Schielzeth 2010, Wolak *et al*. 2012) and have become particularly relevant in the study of labile and repeatedly expressed phenotypes (Bell *et al*. 2009). Coefficients of determination *R*^2^ quantify the variance explained in the fixed part of the model (marginal *R*^2^; *sensu* Nakagawa and Schielzeth 2013). Repeatabilities and coefficients of variation are thus complementary quantities, one focusing on the random the other on the fixed part of the model.

Both, repeatabilities and coefficients of determination, quantify sources of variation in relation to the total variance in a response. To make this more concrete, we want to focus on variance decomposition in a context of the study of phenotypic variation, although the concepts are easily transferred to other systems. Imagine some flexible phenotype of some organism (this may be some physiological, endocrinological or behavioral trait, see e.g. Nespolo and Franco 2007, Bell *et al*. 2009) that has been measured across multiple individuals with repeated observation per individual. Observed phenotypes *y_ij_* are thus clustered within individuals *i* with repeated observations *j* per individual. The phenotypic equation represents a variance decomposition model that consists of a mechanistic (fixed effects) and idiosyncratic (random effects) part (Allegue *et al*. 2017):

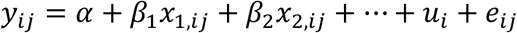

Where *i* indexes individuals, *j* indexes observations, the terms *βx* represent fixed effect predictors *x* and their slopes *β* with numbers indexing different predictors (of which there may be more than the two shown here). The effect of the *βx* terms may be summarized as the linear predictor 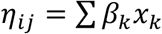 and represents the fixed part of the model. The terms *u* and *e* are the random components, where *u_i_* represents deviations of individuals from the population mean and *e_ij_* represents deviations of observations from individual means. Individual-level deviations *u_i_* and observation-level deviations *e_ij_* are typically assumed to be normally distributed with mean of zero and variance estimated from the data. Since the linear predictor also explains some phenotypic variance, there are three variance components, 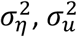 and 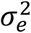, that can be interpreted as sources of biological variation.

Various options for estimating ratios of these variance components for the random and the fixed part of the model have been discussed extensively (O’Dea *et al*. 2021). Here we are concerned with another element, random-slope variation that blurs the distinction between the random and the fixed part of the model. Random slopes are interactions between fixed-effect covariates and random-effects: Slopes that vary by random-effect level (Gelman and Hill 2007, Dingemanse and Dochtermann 2013). Random-slopes are often important, because they allow to control for pseudoreplication in the estimation of the population slope (Schielzeth and Forstmeier 2009, Gurka *et al*. 2011). Furthermore, they represent phenotypically plastic responses to an environment and are therefore relevant in the study of reaction norms (Nussey *et al*. 2007, Dingemanse *et al*. 2010). Random-slope models have become popular in the study of ecology and evolution, because they reflect phenotypic plasticity such as an organisms’ ability to cope with environmental changes (Fox *et al*. 2019).

Random-slope variation disrupts the concept of “repeatability” (Biro and Stamps 2015). With random-slopes, the amount of between-individual variation, for example, depends on levels of the covariate, thus on context, environment, age or time. We have therefore introduced the term “conditional repeatability” for repeatabilities that vary by covariates (Nakagawa and Schielzeth 2010). Although random-slope models have become popular, we are not aware of any universal standardized measure of random-slope variation that is in frequent use. A general framework for variance decomposition in mixed-models has recently been proposed in psychology (Rights and Sterba 2019, 2020). We here focus on particular elements of this framework that we think are most applicable in ecology and evolution. Since meta-analyses on the magnitude of random-slope variation are missing, we also know very little about the magnitude of random-slope variation in natural systems. It therefore needs a system for estimating random-slope variation in a manner that is comparable across study system: It needs a method of standardization.

Johnson (2014) has introduced equations to estimate repeatabilities form random slope models. Johnson’s method is based on the multiplication of the random-intercept random-slope variance-covariance matrix **Σ** with the design matrix **X** for the fixed effects (called **Z** in Johnson 2014). Effectively, Johnson’s repeatability estimates average repeatabilities across the range of the covariate. As such, Johnston’s repeatability is different from and typically larger than then random-intercept variation estimated in the model. In other words, with random-slope variation, the random-intercept variation is no longer a comprehensive parameter that describes the magnitude of individual differences (or differences among other types of groups). Random-intercept variation merely describes individual-differences at a single point of the environment.

Rights and Sterba (2019, 2020) have introduced a comprehensive system for calculating various *R*^2^ measures from mixed-effects models. Unlike Johnson (2014), they use the variances and covariances of fixed effects rather than the design matrices for the estimation. This offers a more concise version for reporting and it is therefore the approach that we adopt below. However, although Rights and Sterba (2019, 2020) strongly argue for single-source *R*^2^ as the main focus of reporting and interpretation, we think that this does not capture the most biologically relevant estimates. An inherent feature of reaction norms that vary across contexts is that the between-individual variance is variable across contexts and an additive decomposition (as shown in Rights and Sterba 2019, 2020) is no longer most efficient.

We here discuss the concept of conditional repeatabilities in some details. We start with theoretical considerations and then proceed to application. In particular, we present a calculation for variance-standardized random-slope variances and make recommendations for comprehensive reporting. Some of the quantities have been proposed by Rights and Sterba (2019), but their value for variance decomposition in ecology and evolution has not yet been appreciated. One important message of our paper is that the variance in intercepts as estimated in random-slope models is a rather arbitrary value (the random effect variance at the point where the covariate is zero), but that a biologically more relevant quantity, the average between-group variance across a meaningful range of the covariate can be easily calculated.

## Theory

### Single random slope with mean-centered covariates

We first consider a phenotypic equation expressed as a mixed effect regression model and assume that all parameters are known with certainty.

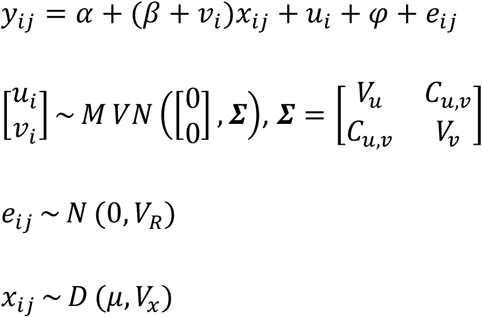

Where *y_ij_* is the response of interest, *α* is the global intercept (an estimate of the population mean if covariates are centered), *x_ij_* is a covariate (context, environment, age or time) that varies within individuals (an observation-level predictor), *β* is the population mean response to the covariate, *u_i_* is the deviation of mean individual trait values from the population mean, *v_i_* is the deviation of individual slopes from the population mean slope and *e_ij_* are residual deviations. *φ* is merely introduced as a placeholder for other additive variance components such as additional fixed or random effects. The variance explained by *φ* is *V_φ_*. The residual deviations *e_ij_* are normally distributed with a variance *V_R_*, and *u_i_* as well as *v_i_* are multivariate normal distributed with variances of *V_u_* and *V_v_*, respectively, and a covariance of *C_u,v_*. The covariate *x_ij_* is arbitrarily distributed (as symbolized by *D*) with mean *μ* and variance of *V_x_*. We will first assume that covariances are grand-mean centered, such that *μ* = 0 and lift the constraint later.

The equation translates into the following variance components (Snijders and Bosker 2011, Allegue *et al*. 2017, Rights and Sterba 2019):

Variance explained by fixed effects: *V_F_* = *β*^2^*V_x_*
Variance explained by individual: *V_I_* = *V_u_* + *V_v_V_x_*
Other variance components: *V_φ_*
Residual variance: *V_R_*
Phenotypic variance: *V_P_* = *V_F_* + *V_I_* + *V_φ_* + *V_R_*

*V_I_* refers to the total variance explained by individual identity, including random intercept and random slope variance. This is an interesting quantity that summarizes individual differences, an important topic of current research (Réale *et al*. 2007, Stamps 2016), but it is different from (generally larger than) *V_u_* if random-slope variation differs from zero. We also call *V_I_* the marginalized variance explained by individual, because it averages (marginalizes) across the range of the covariate.

We note that the phenotypic variance *V_P_* as we calculate it here as the sum of additive variance components might differ slightly from the variance in response values as estimated from the raw data (Rights and Sterba 2020). The difference is that the sum of the variance components aims to estimate the population variance while the variance in raw response values represents to variance in the sample. Since the population variance is what is relevant to biological interpretation (de Villemereuil *et al*. 2018), the sum of additive components is usually preferable. However, if components are not fully additive, this may lead to misestimation. It is therefore important to specify the variance decomposition model correctly.

One component of the total between-individual variation *V_I_* is the variance uniquely explained by random slopes. This is the phenotypic variation explained by individual differences in reaction norms:

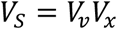

Even though *V_S_* is contributing to *V_I_*, there is no decomposition of *V_I_* in additive components (see below).

### Single random slope with uncentered covariates

We have so far assumed that *μ* = 0, which can be easily achieved by mean-centering covariates prior to the analysis (note that with z-transformation of *x*, *V_S_* = *V_v_*). Covariate centering is generally advisable when used in random-slope models, because uncentered covariates tend to produce large covariation between random-slopes and random-intercepts, which often leads to convergence problems in model fitting. If the covariate was not centered, then

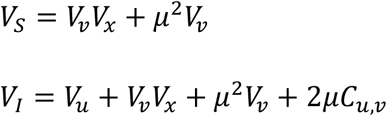

The between-individual variation in average phenotypes is a little more difficult, because as a conditional repeatability it varies across the range of the covariate. We can calculate the amount of between-individual variation for any point *x* as (Figure 1):

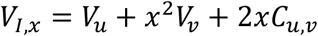

**Figure 1.**
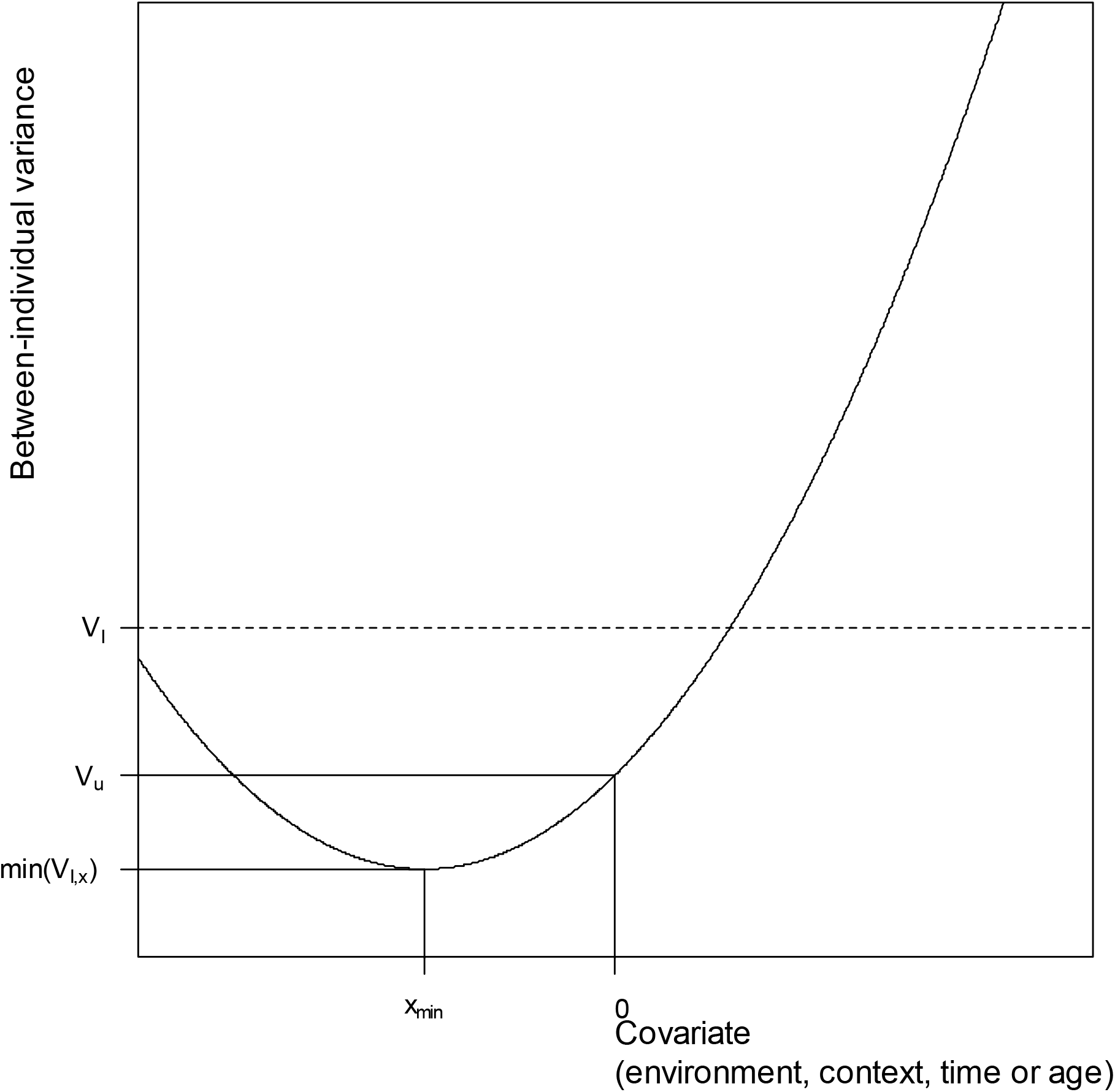
Conceptual display of conditional repeatabilities. The figure shows the between individual variance as it depends on the value of the covariate. The random-intercept variance *V_u_* is always estimated at a covariate value of zero. The minimum between-individual variance *min*(*V_I,x_*) is reached at *x_min_*. The average between-individual variance is *V_I_* and is usually larger than *V_u_* if there is random-slope variation.

The minimum value of *V_I,x_* (from where it increases in either direction) is reached at:

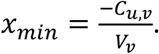

It follows that the minimum value of *V_I,x_* is:

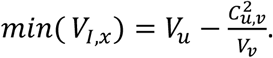

With context-sensitive responses, it is difficult to conceptualize a pure among-individual variation in elevation across the entire gradient (that is, there is no additive decomposition of *V_I_*). We think that *V_I_* is still the best descriptor of overall magnitude in individual differences (Figure 1). One might be tempted to use *V_I*_* = *V_I_* – *V_S_* = *V_u_* as an estimator of elevation, but this is just the between-individual variance at the point where the covariate is zero. Whether this is a meaningful value, depends on how the covariate is centered. The value might be representative for an average covariate value with mean-centered covariates. However, whether or not this is also the minimum value of between-individual variation depends on the intercept-slope covariance that, as a property of the population, is usually beyond experimental control.

### Multiple and correlated predictors

We have above shown equations that use means and variances of covariate *x* to quantify conditional repeatabilities. Johnson (2014) had introduced a different approach that uses the specific design matrix **X** for the fixed effects instead. This approach will be particularly useful when fixed effect predictors are correlated, since positive correlations will inflate the contribution of a predictor to the phenotypic variance beyond *β*^2^*V_x_*.

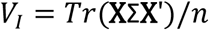

Where **X** is the model matrix for the intercept and the fixed effect of interest (typically a column of 1s for the intercept and columns of covariate values *x* for all observations), **X**′ is the transpose of **X**, *n* is the total number of observations and **Σ** is the random intercept-slope covariance matrix as defined above and as estimated from the data, *Tr* signifies the trace (the sum of the diagonal elements) of the resultant square matrix. To put it simply, *V_I_* is calculated as the predicted amount of between-individual variation associated with all observations and averaged across the dataset.

Johnson’s (2014) approach can be useful for computation, but the alternative use of means and variances of covariates makes reporting much easier. Our simulations (see below) show that means and variances are no less accurate, and yield unbiased estimates in most cases. Furthermore, it is also possible to estimate the variance in the fixed part explained by correlated predictors as:

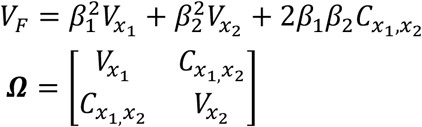

for a model with two fixed effect predictors *x*_1_ and *x*_2_ with variances of *V*_*x*_1__ and *V*_*x*_2__, respectively and predictor covariance of *C*_*x*_1_,*x*_2__. The terms *β*_1_ and *β*_2_ represent the respective population-level slopes. The correlation among predictors is thus summarized in the predictor variance-covariance matrix **Ω**. The predictor variance-covariance matrix **Ω** is suitable for concise reporting and we suggest that it can be reported as part of the supplementary materials with experimental studies.

### Multiple random slopes with uncentered covariates

We have so far assumed that *φ* is an arbitrary, but additive effect. This might include other random-intercept components or additional fixed effects, which are easy to deal with. In general, fixed and random-effects might vary between individuals, such as any group-level fixed effect (like the size of an individual) or higher-level random effect (like the family or patch that an individual originates from). The equations for calculating *V_I_* thus apply to the random-effect variances associated with an individual identity random effect only, that is, after controlling for other between-individual effects in the model. This is generally the case for repeatabilities and might thus be quite familiar.

A different complication arises when additional fixed effects are also involved in random slopes that vary between individuals. This makes the calculations more tedious, but it is still possible to calculate the total variance explained by individual identity across multiple covariates. For example, if there are two covariates *x*_1_ and *x*_2_ that are involved in an interaction with the individual random-effect component to yield random-slopes *v*_1,*i*_ and *v*_2,*i*_.

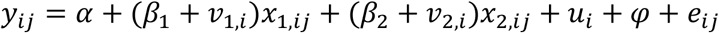

If the means of the two covariates *x*_1_ and *x*_2_ are *μ*_1_ and *μ*_2_, respectively, and the variances in covariates are *V*_*x*_1__ and *V*_*x*_2__, respectively, and the covariance between *x*_1_ and *x*_2_ is *C*_*x*_1_*x*_2__, than the full variance-covariance structure is **Ω** as above. Note that one of the covariates might be the square of the other 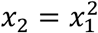, such that all calculations can be used for polynomial model fits, too.

The covariance structure for the random intercepts and slopes is:

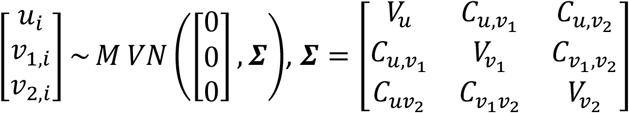

Where *V_u_* represents the random-intercept variance, *V*_*v*_1__ and *V*_*v*_2__ are the random-slope variances for covariates *x*_1_ and *x*_2_ and covariances are shown in the off-diagonal. The total variance explained by the individual random effect can then be calculated as:

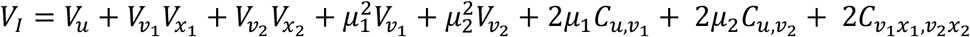

The covariance between the products of random numbers can be calculated as *cov*(*AC, BD*) = E(*A*)E(*B*)*cov*(*C, D*) + E(*C*)E(*D*)cov(*A, B*) + *cov*(*A, B*)cov(*C, D*). In the case of random slopes that are estimated as deviations form the mean, the expectation for *v*_1,*i*_ and *v*_2,*i*_ is zero [E(*v*_1_) = E(*v*_2_) = 0], such that *C*_*v*_1_*x*_1_,*v*_2_*x*_2__ = *μ*_1_*μ*_2_*C*_*v*_1_,*v*_2__ + *C*_*v*_1_,*v*_2__*C*_*x*_1_,*x*_2__. Therefore, the full equation for variance explained by the individual random effect is:

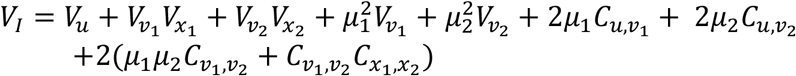

Which can be rearranged to:

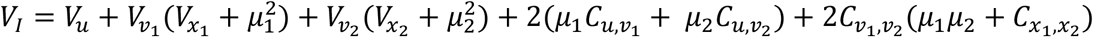

While these calculations seem long and tedious, they can be easily implemented in scripted analysis even for complex models.

## Standardization of variances

The above equations offer obvious ways of variance-standardization for both total between-individual variation *V_I_* and random-slope variation *V_S_*. We propose to call the variance-standardized average between-individual variation *V_I_* the marginalized repeatability *R_mar_*, because it marginalizes (averages) across the environmental gradient:

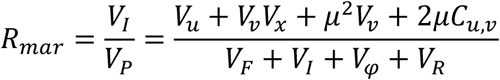

This value is typically larger than the variance-standardized random-intercept variation. We propose to call the variance-standardized random-slope variation the random-slope coefficient of determination 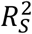 (called 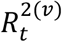 in Rights and Sterba 2019).

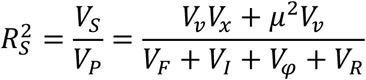

Variance-standardization puts the variance explained by individual components in perspective of the total phenotypic variance, which, in our experience, is what ecologists and evolutionary biologists are usually interested in (see de Villemereuil *et al*. 2018 for a discussion). However, it has been argued that variance-standardization may produce different values not because of differences in the numerator, but because of differences in the denominator (Houle 1992). An alternative way of standardization is therefore standardization by the square of the mean trait value, if the trait is ratio-scale and the assumption that the variance explained increases with the square of the mean is reasonable (Houle *et al*. 2011). For mean-standardization, *V_P_* has to be replaced by 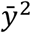 (or *α*^2^ if all covariates are mean-centered) in the equations above.

## Reporting

We write this article partly to encourage complete reporting for future meta-analyses of phenotypic plasticity. A first important message is that the mean of and variance in the covariate are important quantities that allow putting reaction norms and reaction norm variation in perspective of the phenotypic variance. One way to standardize random slopes is the use of variance-standardized covariates 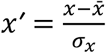, in which case 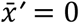 and *V_X′_* = 1. Alternatively, or better additionally, raw mean 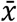 and variance *V_x_* should be reported. Furthermore, it is important to report covariation among fixed effect, ideally in the form of a predictor variance-covariance matrix **Ω**. A second important message is that the correlation among random slopes and random intercepts is an important parameter that can be biologically interpreted and should be reported. Negative correlations show that between-individual variation is lower at high values of the covariate and *vice versa* for positive correlations.

For meta-analysis, it would also need some measures of uncertainty in all relevant estimates. This could be easily achieved by applying our equations to samples from the posterior distributions of models fit in a Bayesian framework (Gelman *et al*. 2004). In a likelihood framework, it could be achieved by parametric bootstrapping (Faraway 2014). However, some of the sampling variances are small in comparison to others and might not need to be available with full uncertainty estimates. For example, estimates of covariate means and variances are estimated with far higher precision than estimates of random effect variances. In fact, it might sometimes be useful to use means and variances for environmental covariates from independent data if the data were collected in an experimental setting where variance in the covariate was manipulated.

## Simulations

We have implemented simple simulation to illustrate some critical point. We do not aim to present a full exploration of the full parameter space, that is potentially vast and varies between applications. For simulations that explore the power of different sampling designs for estimating for estimating random-slope variation we refer to Martin *et al*. (2011), van de Pol (2012) or Westneat *et al*. (2020).

We implemented a data-generating function for a simple random-slope model with a single grouping factor and two covariates. Random-slopes act on an observation-level predictor *x*. Furthermore, a group-level covariate *φ* was introduced with an associated slope *γ*. Random-slopes and random-intercepts were generated from a multivariate normal distribution with means of zero and variance-covariance matrix **Σ**. The generating phenotypic equation as used in our simulations was:

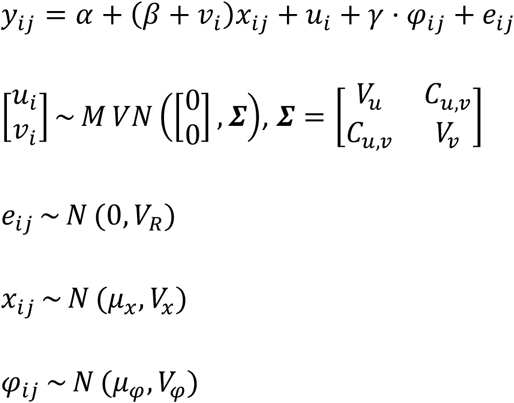

We used the following parameter settings: *α* = 3, *β* = −0.5, *γ* = 0.5, *V_R_* = 1, *V_u_* = 1, *V_v_* = 0.5, *C_u,v_* = 0.3, *V_x_* = 1.2, *V_φ_* = 1, *μ_x_* = 0.5, *μ_φ_* = −0.5. These values were relatively arbitrarily chosen, and effects are purposefully rather strong in order to demonstrate general patterns. The broad patterns are largely insensitive to the detailed choice of values. Covariate values *x* were drawn from uniform distributions. We simulated a population of 60 individuals with an average of 10 observations per individual, which we consider a moderate sample size. For each of 500 simulation runs we fitted the regression model, estimated the parameters and used the above equations to quantify important parts of conditional repeatability. Data were generated in R 4.1.2 (R Core Team 2021) and analyzed using random-slope models fitted in lme4 1.1-27.1 (Bates *et al*. 2015).

The basic setting showed that the conditional repeatability was quite accurately estimated, with only minor bias in *x_min_* and marginal downward bias in *V_I_* (Figure 2a). The accuracy in the estimation of model parameters and derived quantities is unequally distributed with some (like *α, μ_x_, V_x_* and *V_R_*) being estimated with high accuracy while others (like *C_u,v_, V_F_*, *V_φ_* and *x_min_*) being estimated with much less accurately (Figure 3). We also simulated a reduced sample size of 30 individuals and an average of 3 observations per individual. The small-sample simulation resulted substantially increased variation in estimates, but no strong bias except for pronounced bias in *x_min_* (Figure 2b).

**Figure 2.**
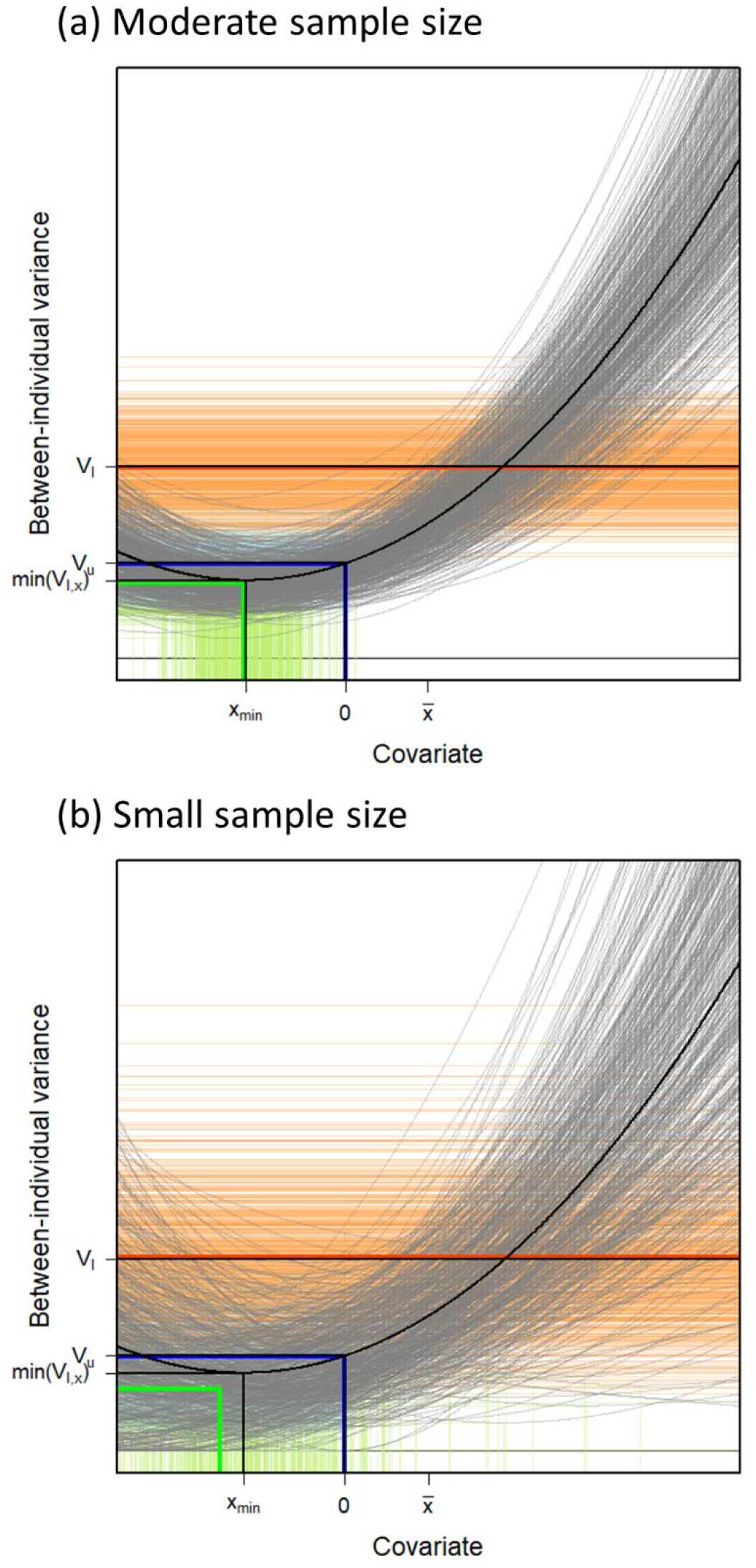
Estimation error in conditional repeatabilities with (a) moderate (*N_ind_* = 60, *N_obs_* = 600) and (b) small (*N_ind_* = 30, *N_obs_* = 90) sample size. The figure shows the same quantities as Figure 1 with the data-generating (true) values shown black. Predicted conditional repeatabilities from 500 replications are shown in grey. Estimated *V_I_*, *V_u_* and *x_min_*/*min*(*V_I,x_*) are shown in orange, blue and green, respectively, with thin lines representing single iterations and bold lines average values.

**Figure 3.**
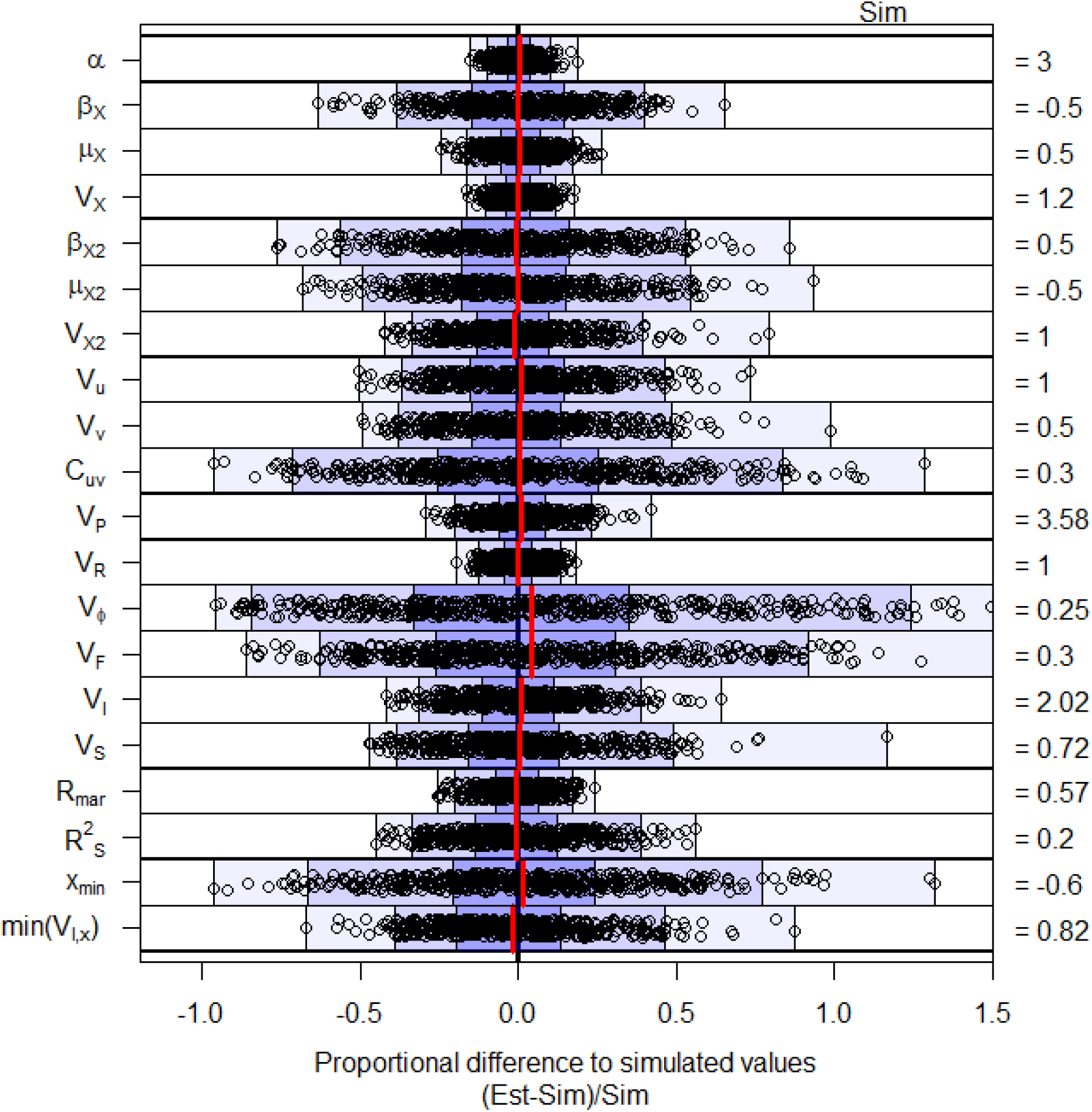
Proportional bias in estimates for various model estimates and derived parameters for a base simulation setting (see main text) with 500 replicates. Results are broadly similar for a range of parameter values when the analysis equation matches with the data generation process. The interquartile range is shown in dark shading, the 95% envelope in moderate shading and the total range in light shading. Dots represent individual estimates from 500 replicates and red lines show mean values. Sim = Expectations based on simulation settings, Est = Estimation based on model fit.

We then assessed estimates when only random intercepts and no random slopes were fitted to the same data (note that data were generated with random-slope variation). These models do not estimate conditional repeatabilities, but rather an overall repeatability. The random-intercept variance was substantially larger than the *V_u_* that was simulated, hence some reaction norm variation in the data was converted to between-individual variation in intercepts in the model. Yet while the estimated was intermediate between *V_u_* and *V_I_*, it substantially underestimated the total individual variance *V_I_* as determined by the phenotypic equation (Figure 4a). Hence, if there is significant reaction norm variation in the population, these models thus typically misestimate average between-individual variation as well as individual variation at the point where the covariate is zero.

**Figure 4.**
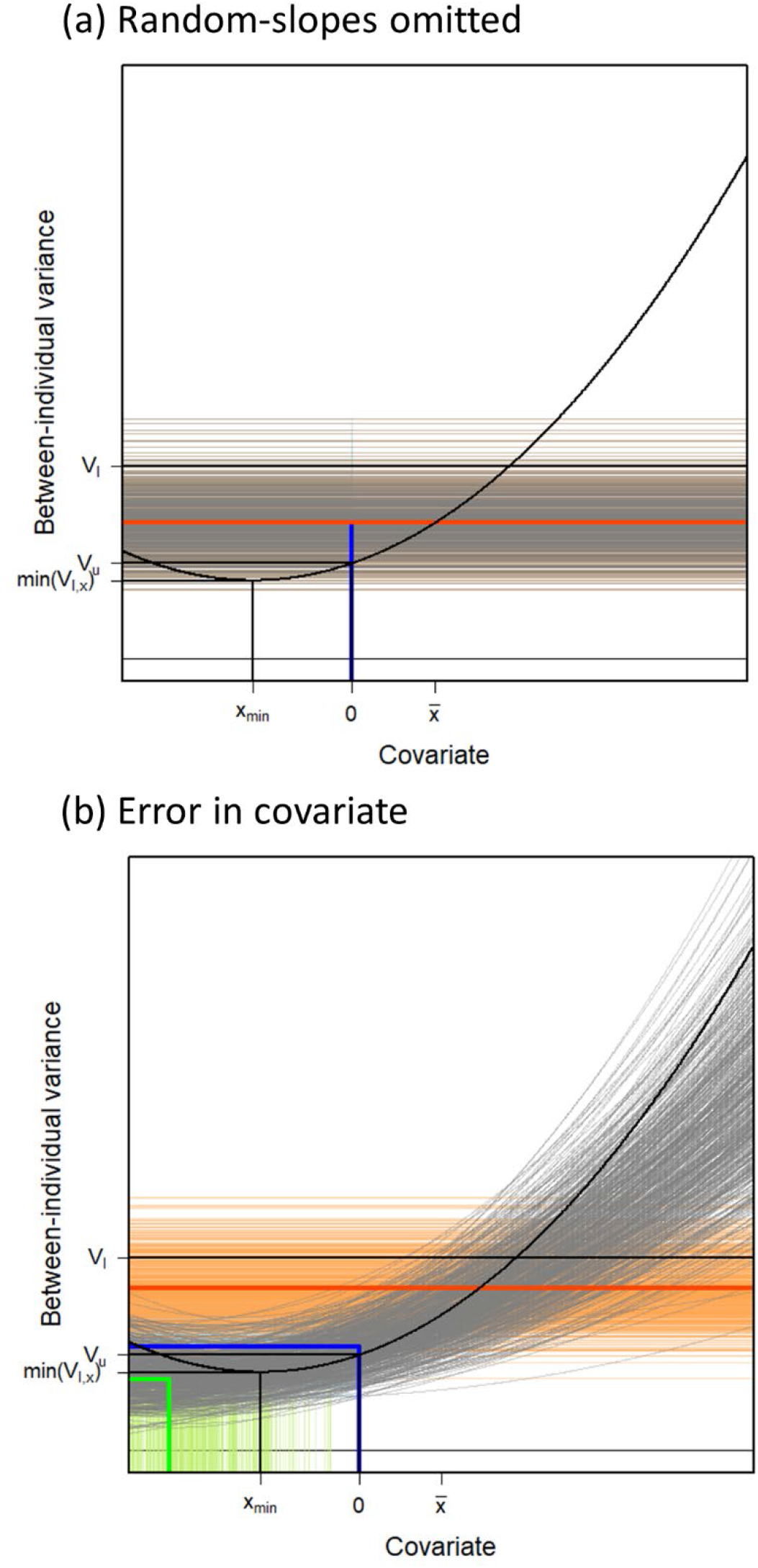
Estimation error in conditional repeatabilities (a) when the analysis model is fitted without random slopes and (b) when there is error in the covariate *x* (30% in this case). The figure shows the same quantities as Figure 1 with the data-generating (true) values shown black. Predicted conditional repeatabilities from 500 replications are shown in grey. Estimated *V_I_*, *V_u_* and *x_min_*/*min*(*V_I,x_*) are shown in orange, blue and green, respectively, with thin lines representing single iterations and bold lines average values. The case of missing random-slopes does not estimate conditional repeatabilities, but the estimated between-individual variance *V_u_* closely approaches the simulated average between-individual variance *V_I_*.

We further explored the effect of measurement error in the covariate on estimates of individual components. Linear models assume that covariates are measured without error (Snijders and Bosker 2011), hence this might sound like a non-sensible attempt. However, measurement error is inevitable, for example when within-subject centering is used to separate within and between-individual responses to some covariate (Raudenbush and Bryk 2002, van de Pol and Wright 2009). Simulations with the same moderate sample size as above and a rather large measurement error of 30% in the covariate resulted in an underestimation of reaction norm variation and underestimation of total between-individual variation *V_I_*, but an overestimation of *V_u_* (Figure 4b). Hence, some of the reaction norm variation was converted to random-intercept variation. This is sometimes unavoidable as in the case of within-subject centering. If the amount of error can be estimated, it could, in principle, be corrected for.

## Conclusions

We present equations that allow the description of conditional between-individual variances (based on Johnson 2014, Rights and Sterba 2019, 2020). Most importantly, we introduce a way of standardized reporting of reaction norm variation, clarify the difference between random-intercept variation and average (marginalized) between individual variation and make recommendations for comprehensive reporting. By putting reaction norm variation in perspective of the phenotypic variance, we aim to promote comprehensive variance decomposition of natural variation in traits of interest. We hope that these tools will stimulate more research on context-sensitivity of individual (or other group-level) variation and will allow data for future meta-analyses to accumulate.

## Acknowledgements

HS were supported by the German Research Foundation (DFG) as part of the SFB TRR 212 (NC^3^) (funding INST 215/543-1, 396782608). SN was supported by the ARC Discovery Project grant (DP180100818).

## Data availability statement

There is no data to be deposited. R scripts for simulations are available on https://github.com/hschielzeth/RandomSlopeR2. The site also holds two R function condR and condRplot that can be used for custom calculations and plotting of conditional variances.

## Author contributions

HS conceived the idea, implemented the simulations and drafted the manuscript; SN helped to refine the method, contributed mathematical derivations and revised the manuscript. All authors gave final approval for publication and agree to be held accountable for the work performed therein.

## Tables

**Table 1:**
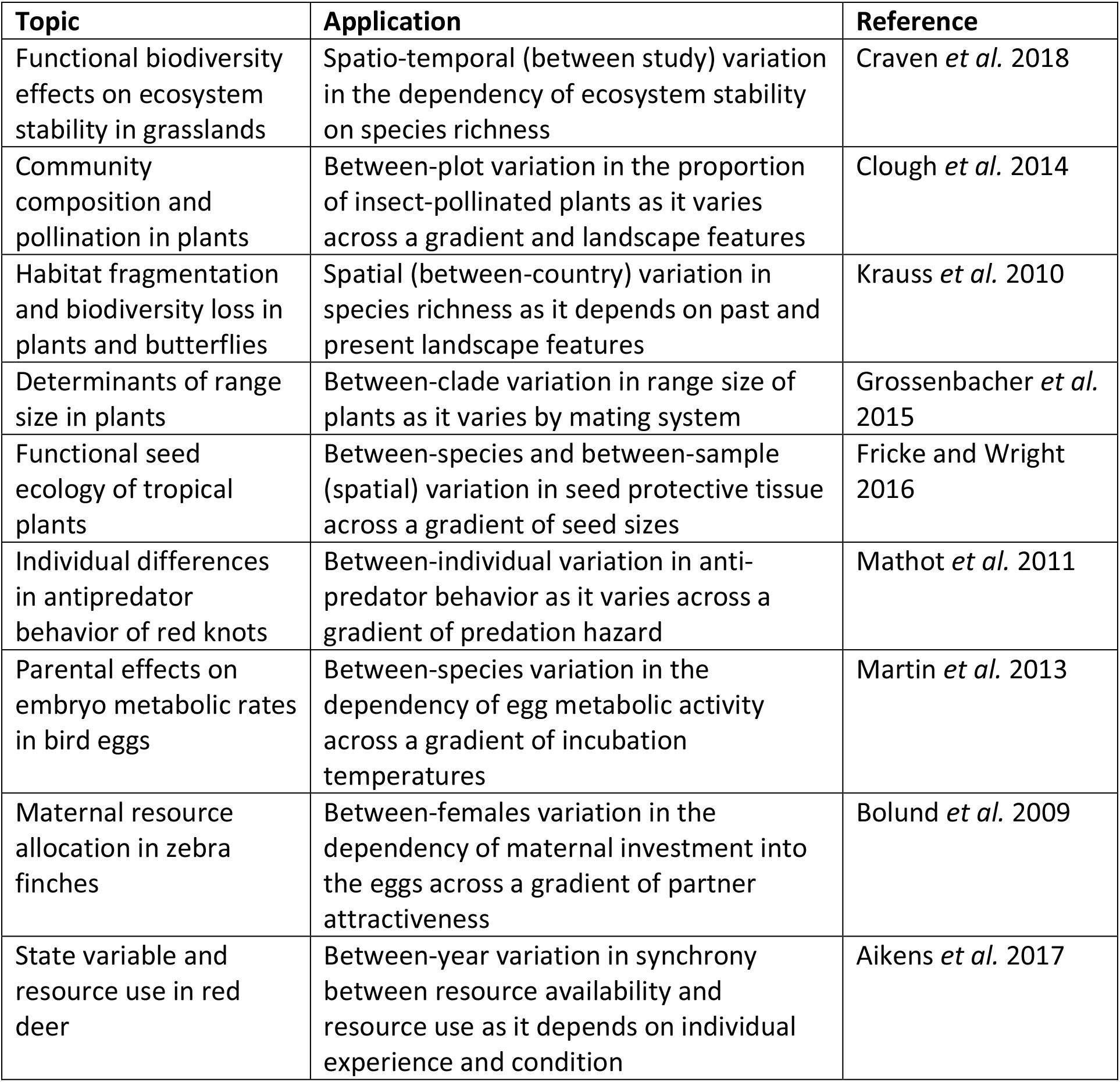
Examples of random slope models in ecological studies with potential application of conditional repeatabilities.

| Topic | Application | Reference |
| --- | --- | --- |
| Functional biodiversity effects on ecosystem stability in grasslands | Spatio-temporal (between study) variation in the dependency of ecosystem stability on species richness | Craven *et al.* 2018 |
| Community composition and pollination in plants | Between-plot variation in the proportion of insect-pollinated plants as it varies across a gradient and landscape features | Clough *et al.* 2014 |
| Habitat fragmentation and biodiversity loss in plants and butterflies | Spatial (between-country) variation in species richness as it depends on past and present landscape features | Krauss *et al.* 2010 |
| Determinants of range size in plants | Between-clade variation in range size of plants as it varies by mating system | Grossenbacher *et al.* 2015 |
| Functional seed ecology of tropical plants | Between-species and between-sample (spatial) variation in seed protective tissue across a gradient of seed sizes | Fricke and Wright 2016 |
| Individual differences in antipredator behavior of red knots | Between-individual variation in anti-predator behavior as it varies across a gradient of predation hazard | Mathot *et al.* 2011 |
| Parental effects on embryo metabolic rates in bird eggs | Between-species variation in the dependency of egg metabolic activity across a gradient of incubation temperatures | Martin *et al.* 2013 |
| Maternal resource allocation in zebra finches | Between-females variation in the dependency of maternal investment into the eggs across a gradient of partner attractiveness | Bolund *et al.* 2009 |
| State variable and resource use in red deer | Between-year variation in synchrony between resource availability and resource use as it depends on individual experience and condition | Aikens *et al.* 2017 |

